# Synthetic Metabolic Pathway for the Production of 1-Alkenes from Lignin-derived Molecules

**DOI:** 10.1101/417857

**Authors:** Jin Luo, Tapio Lehtinen, Elena Efimova, Ville Santala, Suvi Santala

## Abstract

Integration of synthetic metabolic pathways to catabolically diverse chassis provides new opportunities for sustainable production. One attractive scenario is the use of abundant waste material to produce readily collectable product, minimizing production costs. Towards that end, we established the production of semivolatile medium-chain α-olefins from lignin-derived monomers: we constructed 1-undecene synthesis pathway in *Acinetobacter baylyi* ADP1 using ferulate as the sole carbon source. In order to overcome the toxicity of ferulate, we first applied adaptive laboratory evolution, resulting in a highly ferulate-tolerant strain. Next, we demonstrated the 1-undecene production from glucose by heterologously expressing a fatty acid decarboxylase UndA and a thioesterase ‘TesA in the wild type strain. Finally, we constructed the alkene synthesis pathway in the ferulate-tolerant strain. We were able to produce 1-undecene from ferulate and collect the product from the culture headspace without downstream processing. This study demonstrates the potential of bacterial lignin upgradation into value-added products.

## 1 Introduction

The concerns of energy security and environmental issues are driving the development of sustainable and environment-friendly processes for the production of chemicals and fuels. To that end, lignocellulose biorefining has gained substantial attention as a solution to mitigate the dependence on petroleum-based industry (Ragauskas, 2006). Lignocellulose is the most abundant biopolymer on Earth, holding a huge potential as the feedstock for sustainable bioproduction. However, efficient use of lignocellulose is somewhat hindered as lignin, a major component of lignocellulose, is poorly utilized due to its recalcitrance and inherent heterogeneity. Along with the development of second generation biorefineries, an increasing amount of lignin will be generated as a by-product (Ragauskas et al., 2014). In addition, pulp and paper industry produce large quantities of lignin-rich waste as a by-product (Ragauskas et al., 2014; Rinaldi et al., 2016). Thus, it is of high priority to develop technologies for lignin valorization with respect to economic efficiency and environmental sustainability.

To boost the value of lignin, strategies have been developed for lignin depolymerization and subsequent valorization (Abdelaziz et al., 2016). Lignin depolymerization yields various aromatic molecules (monomers, oligomers and polymers), which complicates the downstream processing. Nevertheless, it has been reported that some organisms can utilize these lignin-derived molecules (LDMs) as carbon sources and convert them into central intermediates, protocatechuate and catechol (Abdelaziz et al., 2016; Harwood and Parales, 1996; Linger et al., 2014). The central intermediates will be further converted to acetyl-CoA and succinyl-CoA through β-ketoadipate pathway and enter the citric acid cycle. *Acinetobacter baylyi* ADP1 is one of the microorganisms that has been reported to catabolize various LDMs and even directly depolymerize lignin (Gerischer, 2008; Salmela et al., 2018; Salvachúa et al., 2015). Moreover, *A. baylyi* ADP1 is readily genetically engineered due to its natural transformability and recombination ability (Metzgar et al., 2004), and have been shown to produce a variety of industrially relevant compounds by native and non-native pathways (Lehtinen et al., 2018, 2017, Santala et al., 2014a, 2011). Thus, *A. baylyi* ADP1 could be a potential candidate for lignin valorization.

LDMs are known to be toxic to microorganisms (Cerisy et al., 2017; Ibraheem and Ndimba, 2013; Mills et al., 2009), which also hinders the use of LDMs as substrates. In order to develop a tolerant strain, adaptive laboratory evolution (ALE) can be employed. In ALE, microorganisms are successively cultivated under rationally designed selection pressure, e.g. elevated concentration of inhibitors (Dragosits and Mattanovich, 2013). The approach can lead to the strains with beneficial changes allowing them to adapt to the stressful condition. ALE has been successfully applied on various microorganisms to improve their tolerance against inhibitory compounds (Almario et al., 2013; Atsumi et al., 2010; Cerisy et al., 2017).

Bio-based production of hydrocarbons, such as alkanes and alkenes, is of great interest due to their use as advanced “drop-in” biofuels and various fine chemicals (Choi and Lee, 2013; Lee et al., 2018; Sarria et al., 2017; Zhou et al., 2018). In nature, some organisms have been found to possess the pathways for the synthesis of medium-chain (C8-C12) or long-chain (>C12) hydrocarbons (Sarria et al., 2017; Schirmer et al., 2010). Particularly, medium-chain α-olefins, such as 1-undecene, are attractive molecules for their broad use in detergents, plasticizers and monomers for elastomers (Sarria et al., 2017). In addition, the molecules can accumulate extracellularly and are semivolatile, which allow them to be collected directly from the culture vessel without cell harvesting and/or product extraction, significantly reducing the costs and labour of downstream processing. Recently, a single gene *undA* originated from *Pseudomonas* has been discovered to be responsible for 1-undecene (C11) biosynthesis (Rui et al., 2014). UndA is an oxygen-activating, nonheme iron (II)-dependent decarboxylase that converts free fatty acids (FFAs) to the corresponding terminal alkenes. The enzyme accepts fatty acids with chain length from 10 to 14 as substrates. The gene has been heterologously expressed e.g. in *E. coli* for 1-undecene production (Rui et al., 2014). Other possible alkene synthesis pathways include polyketide synthase (PKS) pathway and head-to-head hydrocarbon synthesis pathway, in which multiple enzymes and reactions are involved (Beller et al., 2010; Liu et al., 2015). In comparison, the one-step decarboxylation of FFAs catalyzed by UndA is simpler and has a narrower substrate spectrum (Kang and Nielsen, 2017).

In this study, we employed *A. baylyi* ADP1 for the production of 1-undecene from a lignin derived model compound, ferulate. We applied ALE to improve the tolerance of *A. baylyi* ADP1 against ferulate, and established a synthetic pathway for the direct conversion of LDMs to 1-undecene (Figure 1). We demonstrate the potential of catabolically diverse bacteria for the synthesis of industrially relevant compounds from an abundant and sustainable substrate.

**Figure 1.**
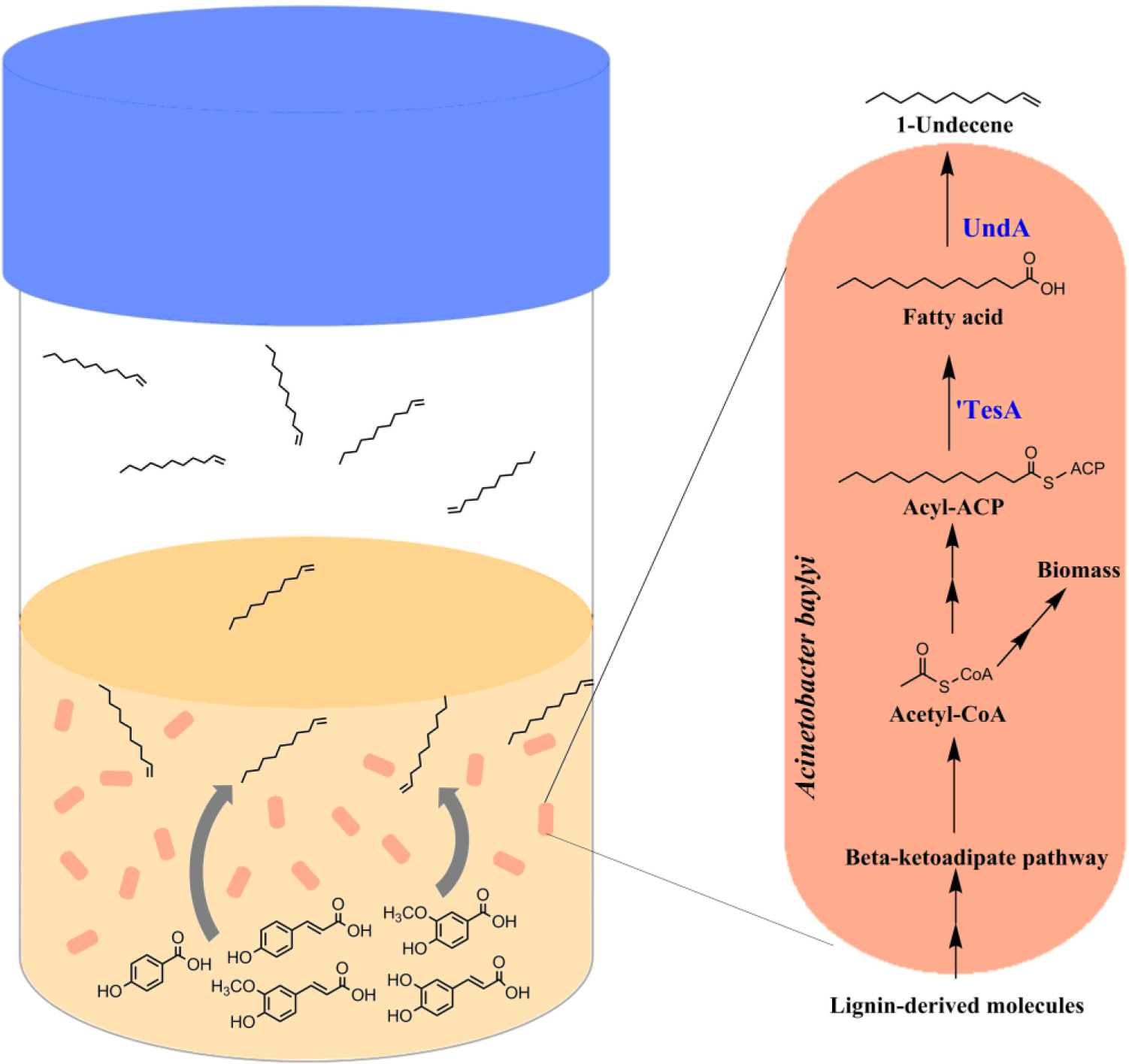
A schematic representation of the medium-chain alkene production from LDMs by *A. baylyi* ADP1. Various LDMs can be converted into central metabolites by *A. baylyi* ADP1. Genes ‘*tesA* and *undA,* encoding for thioesterase and decarboxylase, respectively, are heterologously expressed in *A. baylyi* ADP1 for alkene production. 1-Undecene is a semivolatile hydrocarbon, which can be directly collected from the culture headspace.

## 2 Results

### 2.1 Adaptation of *A. baylyi* ADP1 to high concentration of ferulate

In order to improve the growth of *A. baylyi* ADP1 on the lignin-derived compounds, we carried out ALE using ferulate as the model compound. Ferulic acid is an important building block during lignin biosynthesis (Zhao and Moghadasian, 2008). Its salt form, ferulate, is one of the major lignin-derived aromatic molecules that can be obtained from alkaline pretreated lignin (De Menezes et al., 2017; Vardon et al., 2015). Wild type ADP1 was cultivated in mineral salts medium supplemented with 45 mM ferulate as a sole energy and carbon source, after which the cells were sequentially transferred to fresh medium before reaching a stationary phase. The concentration of ferulate was gradually increased to 125 mM by the end of the evolution, which was the highest concentration introduced. After the evolution, isolates from two concurrently evolved populations were compared. The isolates from population 1 showed better and more consistent growth in ferulate than those from population 2. The best performing isolate, designated as adapted ADP1, was selected for further comparison with wild type ADP1. In adapted ADP1, colony morphology did not change based on the observation on plates. In addition, the natural transformability was maintained after the ALE.

### 2.2 Comparison of growth in ferulate between wild type and adapted ADP1

To compare the growth between wild type and adapted ADP1 in ferulate, both strains were precultivated in mineral salts medium supplemented with 15 mM ferulate, after which cells were transferred to fresh mediums supplemented with different concentrations of ferulate. Generally, adapted ADP1 showed higher tolerance towards ferulate. As ferulate concentration was increased, the growth rates of both strains were reduced (Figure 2A and B), but the degree of reduction for adapted ADP1 was much smaller than that of the wild type. Wild type ADP1 showed poor growth in 80 mM ferulate and the growth was completely inhibited when ferulate concentration was 100 mM. In contrast, adapted ADP1 still exhibited prominent growth in 100 mM ferulate (Figure 2A). In 80 mM, adapted ADP1 also exhibited faster growth than wild type and reached higher optical density (OD) (Figure S2).

**Figure 2.**
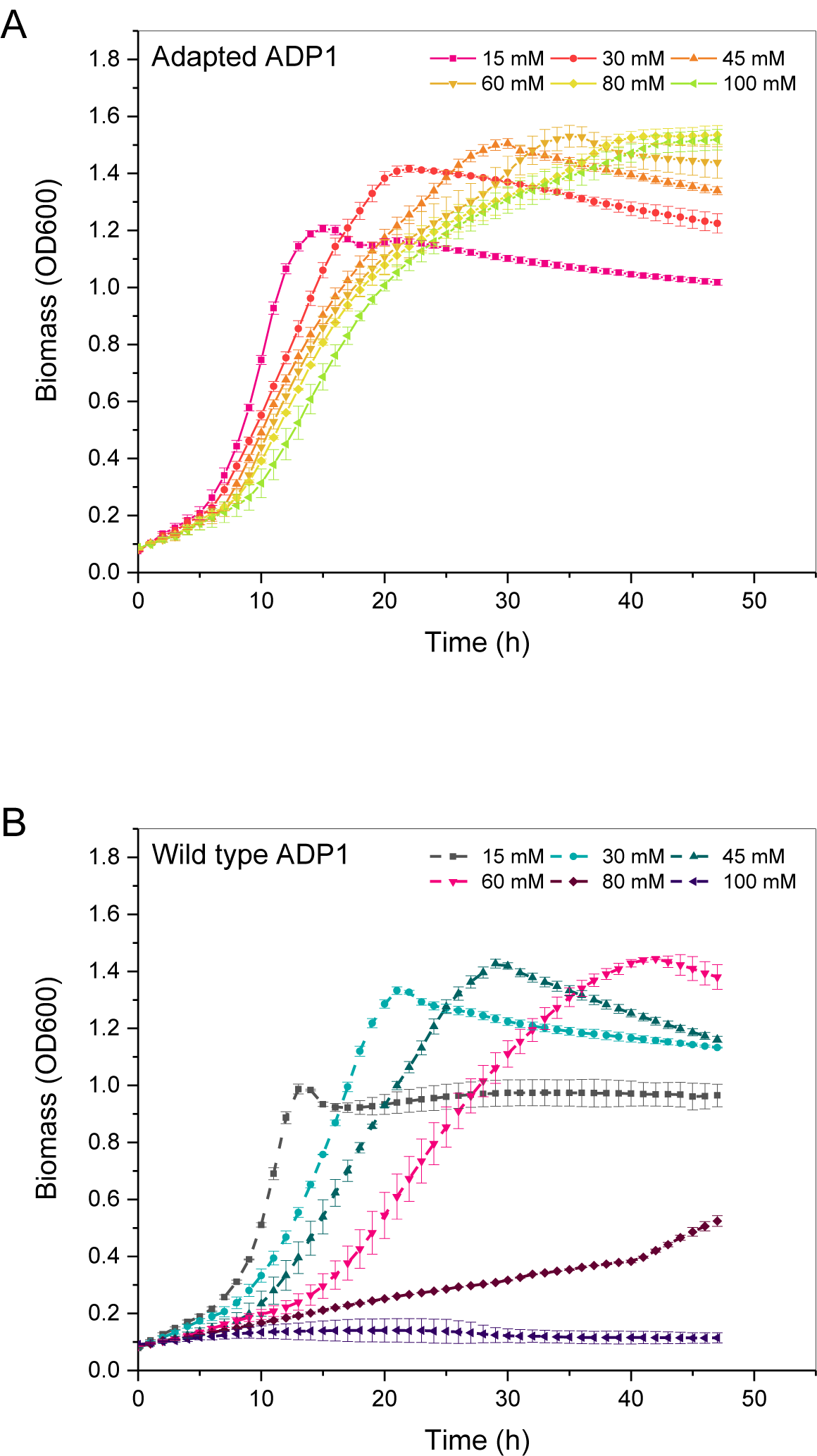
The growth of adapted ADP1 (A) and wild-type ADP1 (B) on ferulate. The strains were cultured in mineral salts medium supplemented with 15 mM, 30 mM, 45 mM, 60 mM, 80 mM, and 100 mM ferulate. The mean values and standard deviations (error bars) from three parallel cultures are shown.

In the aromatic catabolizing pathway of ADP1, ferulate is first converted into vanillate (Fischer et al., 2008). Thus, the growth comparison was also performed using vanillate (15 mM, 45 mM, 75 mM and 100 mM) as a sole carbon source. Similarly, adapted ADP1 showed advantage over wild type in terms of growth on vanillate (Figure S3); Wild type showed slightly slower growth and lower final OD than the adapted ADP1 already in 15 mM vanillate. As the concentration was increased, the growth of wild type ADP1 was inhibited to a larger extent. In comparison, adapted ADP1 exhibited similar growth profile as when grown on ferulate. Interestingly, adapted ADP1 did not show significantly improved tolerance against *p*-coumarate, another LDM which can also be catabolized by ADP1 via β-ketoadipate pathway (data not shown).

### 2.3 Constructing the synthesis pathway for 1-undecene production

To confer 1-undecene production, we heterologously expressed *undA* and *‘tesA* in *A. baylyi* ADP1. The thioesterase ‘TesA (a leaderless version of TesA that is targeted in cytosol), is responsible for the conversion of acyl-ACP to fatty acid, the precursor for alkene synthesis, and therefore hypothesized to benefit the synthesis (Steen et al., 2010). For the expression, a cyclohexanone-inducible promoter ChnR/*P*_*chnB*_ originally isolated from *A. johnsonii* was used (Steigedal and Valla, 2008). The expression system has been previously characterized in *E. coli* and *P. putida* (Benedetti et al., 2016). For testing the functionality of the expression system in *A. baylyi*, a plasmid pBAV1C-chn-GFP was constructed. Induction factor of 52 was obtained for 1 mM cyclohexanone after 3 hours of induction, indicating strong expression and high signal/noise ratio.

Plasmid pBAV1C-chn was constructed and used as the vector for the expression of *undA* and ‘*tesA*. Three different plasmids were constructed: pBAV1C-chn-*undA*, pBAV1C-chn-‘*tesA*, and pBAV1C-chn-‘*tesA-undA* (Figure S1). *A. baylyi* ADP1was then transformed with pBAV1C-chn (empty plasmid control) and the three expression plasmids, designated as ADP1-empty plasmid, ADP1 *UndA*, ADP1 ‘*tesA*, and ADP1 ‘*tesA-undA*.

To select the optimal construct, we compared the production of 1-undecene between the transformants containing different plasmids. The transformants were cultivated in MA/9 medium supplemented with 5% glucose, 0.2% casein amino acid, and 25 μg/ml chloramphenicol. The cells were induced with 1 mM cyclohexanone when the OD reached 1. After 1 h of induction, the cells were transferred to sealed vials and cultivated overnight. The produced and evaporated alkenes were directly analyzed from the headspace of the culture vials by solid phase micro-extraction (SPME)–gas chromatography mass spectrometry (GCMS). The production of 1-undecene by different strains is shown in Figure 3. ADP1-empty plasmid produced only traces of 1-undecene (4.46 ± 0.07 μg/L). The production was greatly increased by expressing *undA* alone in ADP1 (418 ± 24 μg/L) while expressing ‘*tesA* alone did not have a great influence on the production (5.06 ± 0.33 μg/L). ADP1 ‘*tesA-undA* had the highest production (694 ± 76μg/L), which was 1.7 fold higher than with ADP1 *undA*. In addition, signal of 1-tridecene were detected in the cultivation with ADP1 ‘*tesA-undA* (Figure S5). At the end of the cultivation, ADP1 with empty plasmid and ADP1 *undA* reached the OD of more than 2 while the other strains had the OD approximately 1.5 (Figure S4). Based on the results, pBAV1C-chn-‘*tesA-undA* was selected for 1-undecene production from ferulate.

**Figure 3.**
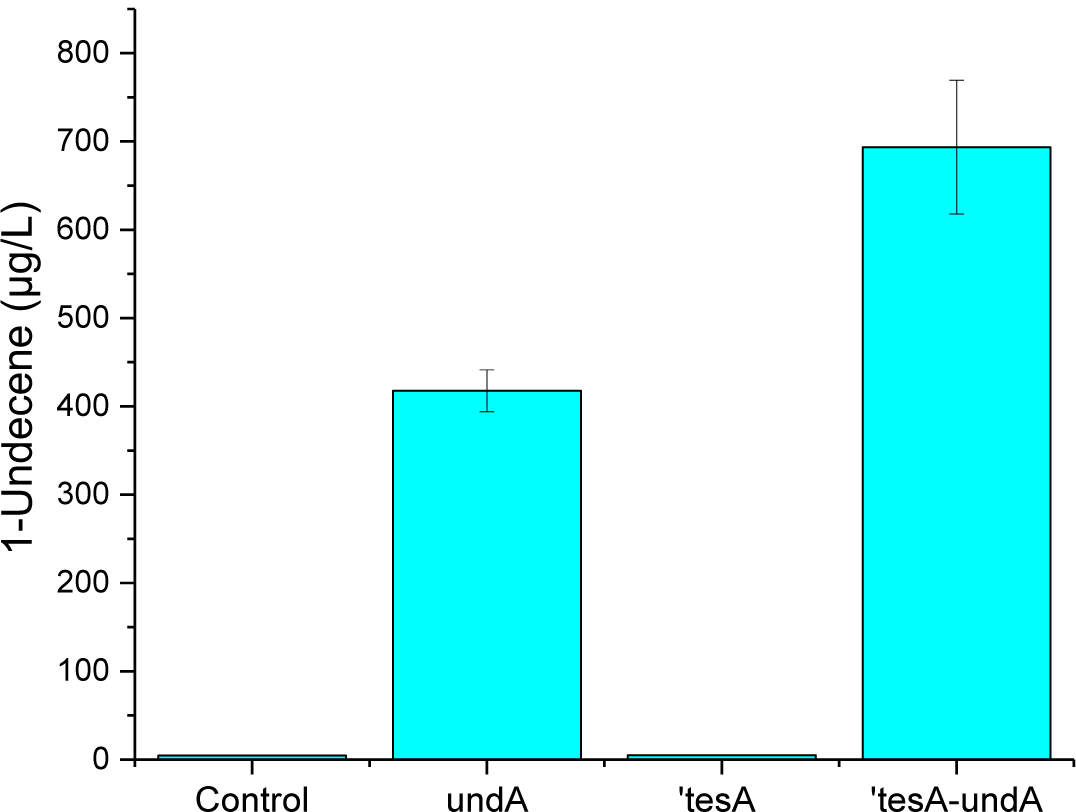
Production of 1-undecene from glucose by *A. baylyi* ADP1 with different constructs. In the histogram, from the left to the right are 1-undecene productions with ADP1-empty plasmid pBAV1C-chn (control), ADP1 *UndA*, ADP1 ‘*tesA*, and ADP1 ‘*tesA-undA*. The mean values and standard deviations (error bars) from two parallel cultures are shown.

#### 2.4 1-Undecene production from ferulate with adapted ADP1

We demonstrated that the highest 1-undecene production was obtained with pBAV1C-chn-‘*tesA-undA* among the constructed plasmids. Thus, adapted ADP1 was transformed with the construct and designated as adapted ADP1 ‘*tesA-undA*. Both adapted ADP1 ‘*tesA-undA* and ADP1 ‘*tesA-undA* were precultivated in mineral salts medium supplemented with 5 mM ferulate, followed by batch cultivations in 110 mM ferulate. During the cultivation, biomass and ferulate concentration were monitored. The detection of 1-undecene was carried out by SPME-GCMS.

As expected, adapted ADP1 ‘*tesA-undA* showed distinct advantage over ADP1 ‘*tesA-undA* when cultivated in approximately 110 mM ferulate (Figure 4 A). ADP1 ‘*tesA-undA* showed almost no growth or ferulate consumption during the cultivation, the final OD being 0.21. On the contrary, adapted ADP1 ‘*tesA-undA* showed efficient growth in approximately 110 mM ferulate; the OD was 3.6 at the time of induction, and 5.5 after the following 4.5 h of incubation with the inducer. At this time-point, 52 mM ferulate was left in the culture.

**Figure 4.**
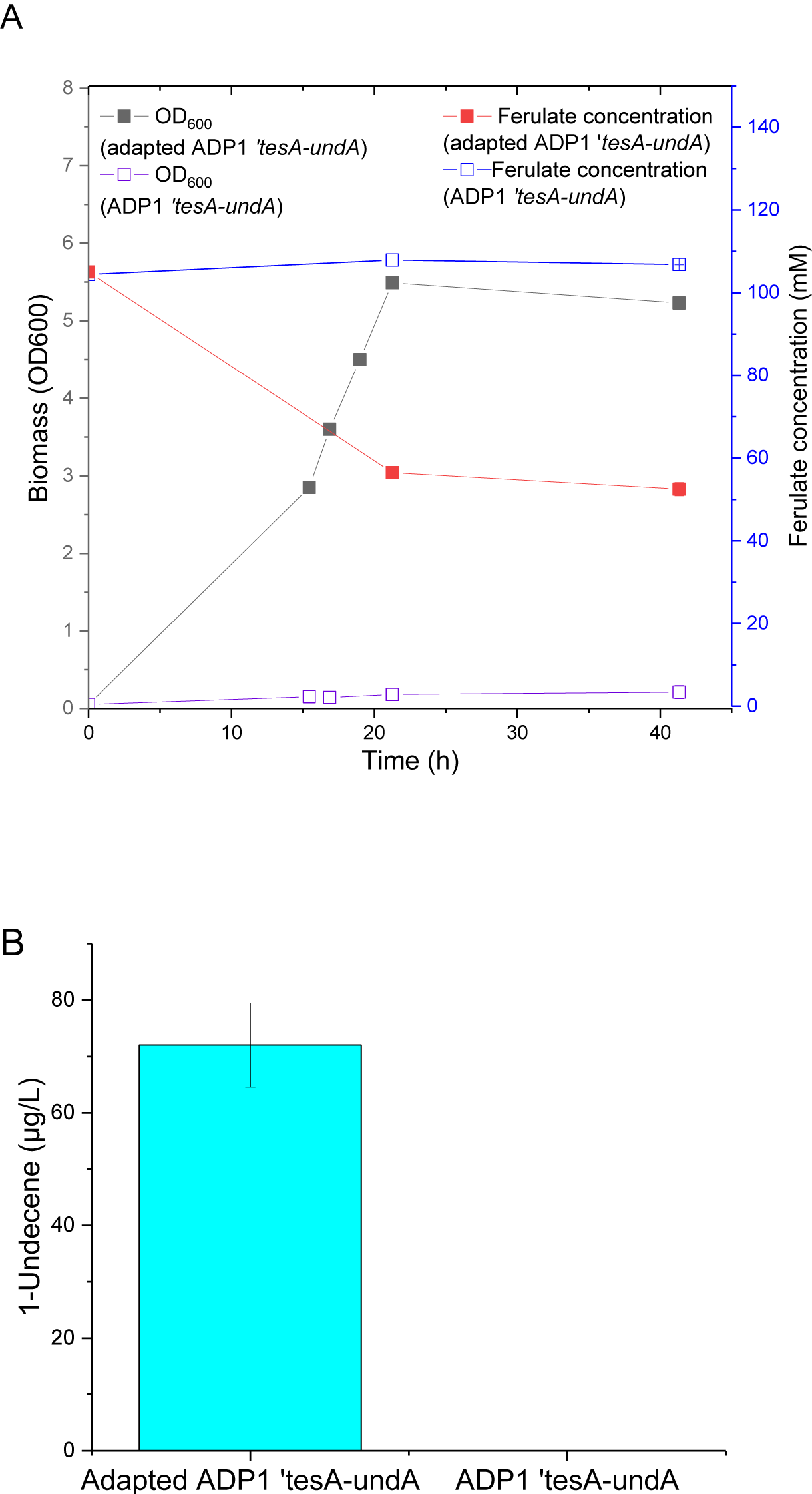
Growth and 1-undecene production by ADP1 *‘tesA-undA* and adapted ADP1 ‘*tesA-undA* from ferulate. (A) The biomasses (OD) and ferulate concentrations of adapted ADP1 ‘*tesA-undA* and ADP1 ‘*tesA-undA.* The strains were cultivated in mineral salts medium supplemented with 110 mM ferulate. The cells were induced after 17 h of incubation in aerated flasks, and thereafter incubated in sealed vials for 20 hours. The values of the last sampling point are the mean from two parallel cultures and the error bars represent the standard deviations (the standard deviations are low, and thus the error bars are not visible in the figure). (B) 1-Undecene production from ferulate as the sole carbon source. 1-undecene was directly collected from the culture headspace. The mean values and standard deviations (error bars) from two parallel cultures are shown.

During the cultivation in sealed vials, due to the oxygen limitation, the OD did not change greatly and only a small amount of additional ferulate was consumed. At the end of cultivation, 72 ± 7.5 μg/L 1-undecene was detected from the cultivation of adapted ADP1 ‘*tesA-undA* while no 1-undecene was produced by ADP1 ‘*tesA-undA* (Figure 4 B).

## 3 Discussion

Lignin has a great potential as a sustainable substrate for bio-based production of fuels and chemicals: It has high energy and carbon content, it is renewable and widely available, and large quantities of lignin-rich waste streams are generated by e.g. bioethanol and paper industry. The practical problem with its utilization is that the typically employed production hosts cannot degrade and further metabolize the aromatic compounds that are the constituents of lignin (Beckham et al., 2016). Furthermore, the aromatic compounds are strong growth inhibitors (Mills et al., 2009). In this study, we established a novel bacterial production platform that utilizes a major constituent of lignin, ferulate, as the sole carbon source. *A. baylyi* ADP1 was employed as the host due to its ability of aromatic compound utilization, adaptability, and the possibility to funnel the intermediates to products of interest.

Although *A. baylyi* ADP1 can utilize ferulate as a sole carbon source, it was found out that the growth rate is reduced or completely inhibited at concentrations relevant for a bioprocess. The mechanism of the inhibition caused by ferulate has not been characterized in detail, but a general mechanism of the inhibition caused by phenolic compounds is thought to be related to their hydrophobicity (Fitzgerald et al., 2004; Mills et al., 2009); phenolic compounds can target cell membranes and interact with lipids and membrane-embedded proteins, breaking the integrity of the membranes. In order to allow the use of LDMs as a substrate for both cell growth and product synthesis, it is a prerequisite to overcome the toxicity of these compounds to host cells. To this end, ALE was employed for *A. baylyi* ADP1 to improve the tolerance and growth on ferulate. As a result of ALE, the growth inhibition caused by ferulate was significantly reduced. The adapted strain showed not only robust growth in high ferulate concentration (up to 125 mM) while the growth of wild type was completely inhibited, but also more prominent growth than wild type in low ferulate concentration. In a previous study, the genes involved in the tolerance towards coumaric acid, another lignin-derived phenolic compound, were identified in *Pseudomonas putida* (Calero et al., 2018). Most of the identified genes were related to membrane stability, transport system, and membrane proteins while the genes involved in the degradation of the compounds only play minor roles in the tolerance (Calero et al., 2018). Those identified genes seem to be more related to global stress handling. In this study, it seems that the improved tolerance is resulted from a different mechanism, since the adapted strain shows improved tolerance specific to ferulate and vanillate but not to coumarate. The additional methoxyl group on the benzene ring of ferulate and vanillate may play an important role in the mechanism. The correlations between the improved phenotype and the genotype remain to be investigated in future.

Alkenes represent industrially relevant platform chemicals used in a broad range of applications and products. Interestingly, the short and medium chain molecules are semivolatile, potentially allowing straight-forward collection and continuous production processes. As a proof-of-concept, our aim was to demonstrate the production of 1-undecene from ferulate without downstream processing. To confer the pathway for alkene production, three different plasmids were constructed to express either undA, ‘tesA, or both undA and ‘tesA in the wild type ADP1. Free fatty acids with chain lengths from 10 to 14 typically serve as the substrate for UndA, whereas acyl-CoA and acyl-ACPs are unlikely converted to alkenes (Rui et al., 2014). However, the strain possessing only *undA* produced 418 ± 24 μg/L 1-undecene (93-fold more than the control strain) from glucose, indicating the existence of available FFAs in ADP1 for 1-undecene synthesis. It has been reported that *A. baylyi* possesses a natural intracellular thioesterase, which can produce FFAs ranging from C6 to C18 from acyl-ACP (Zheng et al., 2012), potentially explaining why UndA alone can confer alkene production in ADP1. The trace amounts of 1-undecene detected in the control strain indicates the existence of natural alkene synthesis mechanism in ADP1. The expression of ‘*tesA* can further convert acyl-ACPs to FFAs, which provides more precursors for 1-undecene synthesis. Although ‘TesA prefers C14 acyl-ACPs as the substrate, it also accepts C12 acyl-ACP, the precursor for 1-undecene synthesis (Choi and Lee, 2013). Hence, it was expectable that ADP1 ‘*tesA-undA* produced the most 1-undecene, 694 ± 76μg/L, improving the production by 1.7 fold in comparison with ADP1 *undA* alone.

Since it was demonstrated that the co-expression of *undA* and ‘*tesA* improved the production of 1-undecene in the ADP1 wild type, the same construct pBAV1C-chn-‘*tesA-undA* was transformed to the adapted ADP1 to produce 1-undecene from ferulate. After the transformation, the adapted strain still maintained excellent growth in high ferulate concentration. 1-Undecene with a concentration of 72 ± 7.5 μg/L was successfully produced, when ferulate was used as the only carbon source. The biomass and ferulate concentration did not significantly change during the cultivation in sealed vials, which is probably due to the limitation of oxygen.

In a previous study, Chen et al. (2015) expressed the one-step decarboxylase OleT in *S. cerevisiae* and increased the production of total intracellular alkenes 67.4-fold to 3.7 mg/L by combinatorial metabolic engineering strategies and process optimization. Liu et al. (2014) expressed OleT in FFAs overproducing *E. coli* and obtained 97.6 mg/L of total intra-and extracellular alkenes, the highest reported production so far. The availability of FFAs plays an important role in alkene production. However, in the aforementioned studies, most alkenes produced had chain lengths of 15 and 17, only small amount of 1-undecene (C11) was produced. Compared to OleT, UndA has a narrower substrate spectrum (C10 to C14). Rui et al. (2014) overexpressed an UndA homolog in *E. coli* and obtained 6 mg/L extracellular 1-undecene production without strain optimization. UndB was later found to be the most efficient among UndA and OleT for 1-undecene production (Rui et al., 2015). Co-expression of Pmen_4370, an UndB homolog, and UcFatB2, a C12-specific thioesterase for lauric acid (C12) synthesis, conferred extracellular 1-undecene production of a titer of ∼55 mg/L in *E. coli* (Rui et al., 2015). Thus, the alkene chain length can be controlled by the specificity of the key enzymes towards the substrates.

Compared to the previous studies, the 1-alkene titers obtained here were low, 694 ± 76 μg/L from glucose and 72 ± 7.5 μg/L from ferulate. However, it should be noted that in the latter case, all the required energy and carbon for both generating the catalyst (biomass) and the production of 1-undecene was obtained from ferulate, emphasizing the potential of the used cell platform. In addition, as the goal was to demonstrate the direct production and collection of 1-undecene without downstream processing, only the alkenes that were secreted by the cells were analyzed. In order to improve the production, the culture set-up should be further developed and optimized; in the current system, the cell growth and consequently 1-undecene production rapidly cease due to the oxygen limitation in sealed vials. For example, cultivations in a bioreactor enabling continuous culture and product recovery would likely significantly improve the productivity. In addition, employing molecular level strategies, such as increasing the availability of FFAs and co-factors, selection of more efficient enzymes specific to medium-chain length substrate, and blocking of competing pathways could further improve the production (Chen et al., 2015; Kang and Nielsen, 2017; Peralta-Yahya et al., 2012).

As demonstrated in this study medium-chain alkenes can be produced directly from ferulate through microbial conversion. However, to realize the upgrading of lignin to medium-chain alkenes, many factors should be considered. For example, efficient substrate conversion is the key factor for high productivity. Technologies for lignin depolymerization, such as alkaline pretreatment of lignocellulose, yields low molecular weight LDMs which can serve as the substrates for microbial conversion (Abdelaziz et al., 2016). However, considering that the depolymerization of lignin gives rise to a highly heterogeneous mixture of different acids and phenolic compounds, it is necessary to evaluate the utilization of mixed LDMs and even real depolymerization products. It has been reported that the catabolism of aromatic compounds via β-ketoadipate pathway is affected by different regulatory mechanisms (Bleichrodt et al., 2010; Fischer et al., 2008). In order to allow efficient utilization of mixed substrates, further engineering should be employed to reduce the carbon catabolite repression, e.g., by deletion of regulatory elements (Vardon et al., 2015). In addition, improvement of the tolerance against mixed LDMs is also crucial for obtaining high substrate conversion rate.

## 4 Conclusions

In this study, we aimed to demonstrate the potential of lignin-derived compounds as substrates for the bioproduction of industrially relevant compounds. To that end, we established the production of α-olefins (namely 1-undecene) by a synthetic pathway in *A. baylyi* ADP1 using ferulate, a model compound for lignin monomer, as the sole carbon source. The tolerance of *A. baylyi* ADP1 against ferulate was significantly improved by ALE, which allowed the use of high ferulate concentration for the production. Our study emphasizes the importance of host selection, and promotes the use of *A. baylyi* ADP1 as a potential chassis for lignin valorization.

## 5 Methods

### 5.1 Strains

*E.coli* XL1-Blue (Stratagene, USA) was used for plasmid construction and amplification. Wild type *A. baylyi* ADP1 (DSM 24193) was used as the parental strain in ALE to develop adapted ADP1. Wild type *A. baylyi* ADP1 was transformed with plasmid pBAV1C-chn, pBAV1C-chn-*undA*, pBAV1C-chn-*‘tesA*, pBAV1C-chn-*undA-‘tesA* and pBAV1C-chn-*‘tesA-undA*, designated as ADP1 empty plasmid, ADP1 *undA*, ADP1 ‘*tesA*, ADP1 *undA*-‘*tesA* and ADP1 ‘*tesA-undA,* respectively. ADP1 empty plasmid was used as control. The adapted ADP1 was transformed with pBAV1C-chn-*‘tesA-undA* and designated as adapted ADP1 ‘*tesA-undA*.

## 5.2 Media

Luria-Bertani medium (10 g/L tryptone, 5 g/L yeast extract, 1 g/L NaCl) was used for plasmid construction and transformation of wild type *A. baylyi*ADP1. The medium was supplemented with 1% glucose as carbon source and 25 μg/ml chloramphenicol as antibiotic when needed. For solid medium, 15 g/L agar was added.

Minimal salts medium (MA/9) was used for the cultivation of plasmid selection. The composition was as follow:Na_2_HPO_4_ 4.40 g/L, KH_2_PO_4_ 3.40 g/L, NH_4_Cl 1.00 g/L, nitrilotriacetic acid 0.008 g/L, NaCl 1.00 g/L, MgSO_4_ 240.70 mg/L, CaCl_2_ 11.10 mg/L, FeCl_3_ 0.50 mg/L. The medium was supplemented with 0.2% casein amino acid, 5% glucose and 25 μg/ml chloramphenicol when indicated.

Mineral salts medium as described by Hartmans et al. (1989) was used for the ALE cultivation, the growth comparison in ferulate (wild type ADP1 vs. adapted ADP1), the transformation of adapted ADP1 and the cultivation of 1-undecene production from ferulate. This medium was applied in the experiments related to adapted ADP1 to maintain its evolved properties, as the strain was adapted with the medium. The composition was shown as follow: K_2_HPO_4_ 3.88 g/L, NaH_2_PO_4_ 1.63 g/L, (NH_4_)_2_SO_4_ 2.00 g/L, MgCl_2_.6H_2_O 0.1 g/L, Ethylenediaminetetraacetic acid (EDTA) 10 mg/L, ZnSO_4_.7H_2_O 2 mg/L, CaCl_2_.2H_2_O, 1 mg/L, FeSO_4_.7H_2_O 5 mg/L, Na_2_MoO_4_.2H_2_O 0.2 mg/L, CuSO_4_.5H_2_O 0.2 mg/L, CoCl_2_.6H_2_O 0.4 mg/L, MnCl_2_.2H_2_O 1 mg/L. This medium was supplemented with different concentrations of ferulate (15-125mM for ALE, 15-100 mM for growth comparison, 100 mM for the transformation of adapted ADP1 and 110 mM for 1-undecene production). To prepare the stock solution (200 mM) of ferulate, proper amount of ferulic acid (Sigma-Aldrich) was added to deionized water, after which equimolar amount of NaOH was slowly added while stirring until ferulic acid was completely dissolved. For solid medium, 15 g/L agar was added. Chloramphenicol (25μg/ml) was added when needed.

### 5.3 Adaptive laboratory evolution

The ferulate-tolerant strains were evolved by a short-term serial passage in mineral salts medium supplemented with ferulate as sole carbon source. Wild type *A. baylyi* ADP1 was first cultivated on solid mineral salts medium containing 15 mM ferulate. Single colony was selected and precultivated in Erlenmeyer flask (100 ml) containing 10 ml medium supplemented with 45 mM ferulate at 30 ^°^C, 300 rpm. When reaching log phase, the cells were cryopreserved as parental strain and the culture was passaged to two Erlenmeyer flasks (100 ml) containing the same medium as in the precultivation. The two populations were evolved in parallel. The concentration of ferulate was initially 45 mM and gradually increased during the evolution to maintain the selection pressure. Cells were passaged to fresh medium before entering into stationary phase to avoid unwanted mutation. Before each passage, the OD of the culture was measured. The amount of inoculum was adjusted daily to make the initial OD of each cultivation between 0.03 and 0.1. The cells were cryopreserved at −80^°^C every two passages. The evolution went on for two months and 61 transfers were performed, corresponding to more than 350 generations. After the evolution, single colonies from both populations were screened out on plates. The strain with the best growth in ferulate, designated as adapted ADP1, was further compared with wild type ADP1 (the parental strain).

## 5.4 Comparison of growth in ferulate between wild type and adapted ADP1

Wild type and adapted ADP1 were precultivated in 5 ml mineral salts medium supplemented with 15 mM ferulate at 30 ^°^C and 300 rpm. After 24 h, the cells from both precultures were transferred to 96-well plate containing 200 μl mineral salts medium supplemented with different concentrations of ferulate (15 mM, 30 mM, 45 mM, 60 mM, 80 mM and 100 mM) respectively. Each cultivation was performed in triplicate. The cells were then incubated in Spark multimode microplate reader (Tecan, Switzerland) at 30 ^°^C for 72 h. Shaking was performed for 5 min twice an hour with a frequency of 54 rpm. The optical density at 600 nm (OD600) was measured every hour.

Wild type and adapted ADP1 were also compared regarding the growth in vanillate, the metabolite derived from ferulate in the aromatic catabolizing pathway. The same processes as above were used for the comparison but vanillate was used instead of ferulate.

## 5.5 Plasmid construction and transformation

Plasmid construction was carried out using *E. coli* XL-1 Blue as host. The reagents for PCR, digestion and ligation were provided by Thermo Scientific (USA) and used according to provider’s instruction. The primers used in this study are listed in table 1.

**Table 1.**
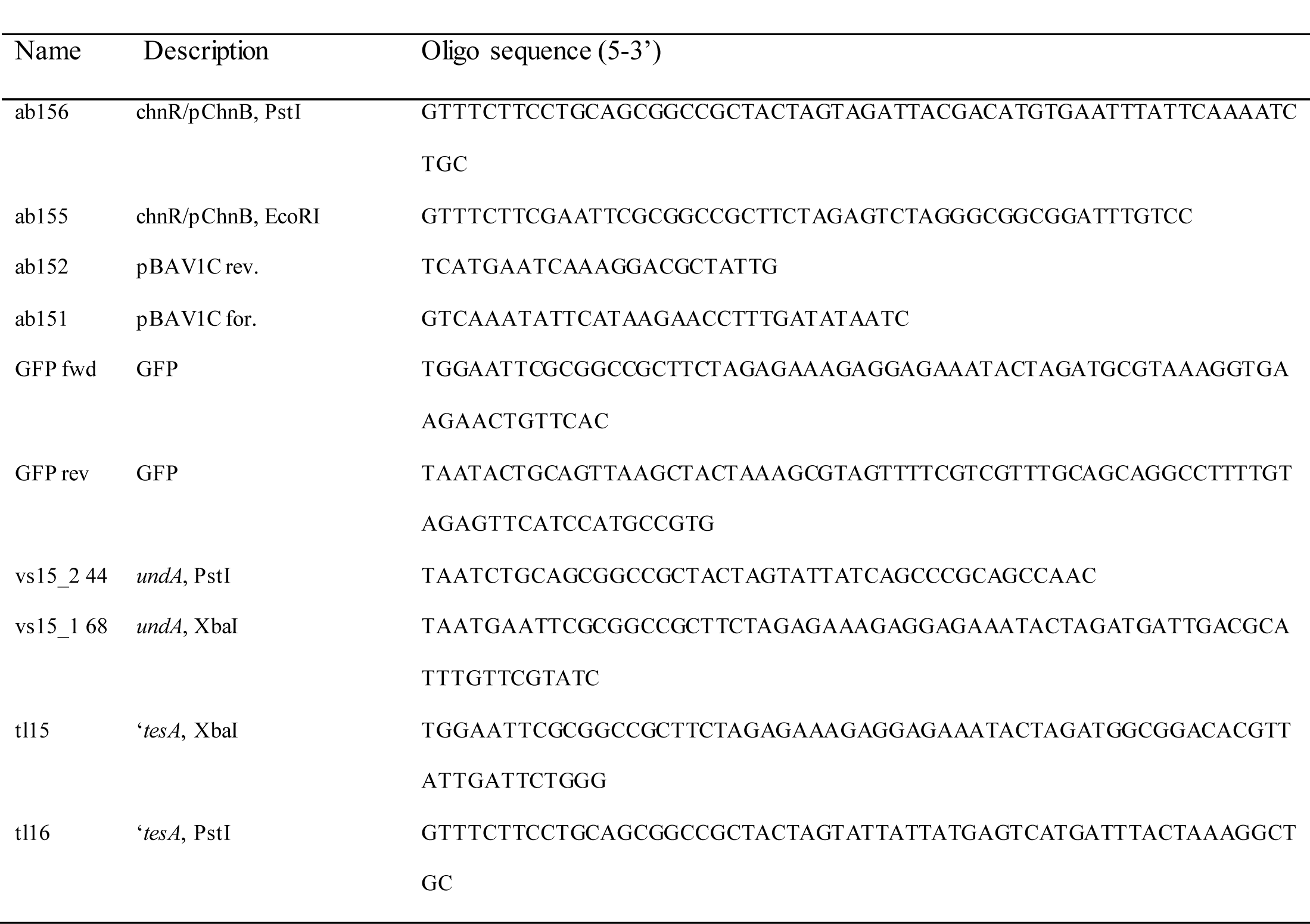
List of primers used in this study

The plasmid pBAV1C-chnR/pChnB designated as pBAV1C-chn was constructed as follows. First, the plasmid pBAV1C-T5-GFP, which was constructed by Santala et al. (2014b), was amplified with the primers ab151 and ab152 to remove internal NdeI-site from the plasmid backbone. The PCR product was self-ligated and transformed to *E. coli* XL1-Blue strain. The obtained plasmid was named as pBAV1C-T5-GFP. Then, the fragment containing the regulator chnR and its cognate promoter pChnB was amplified from plasmid pSCM (a kind gift from Standard European Vector Architecture database (Spain)) (Benedetti et al., 2016), using primers ab155 and ab156. To obtain pBAV1C-chn, the amplified fragment was cloned to the pBAV1Cd plasmid backbone according to the BioBrick assembly standard 10 using EcoRI and PstI sites.

For the initial characterization of cyclohexanone-inducible promoter system (*ChnR*/*PchnB*) in *A. baylyi* ADP1, a plasmid pBAV1C-chn-GFP was constructed as follows. The gene fragment encoding monomeric and superfolder green fluorescent protein variant (GFP) was PCR amplified from the pSCM plasmid using biobrick suffix and prefix primers, GFP fwd and GFP rev. The fragment was BioBrick-cloned to pBAV1C-chn plasmid to obtain pBAV1C-chn-GFP plasmid which was then transformed into *E. coli* XL-1 Blue. Verified pBAV1C-chn-GFP construct was transformed into *A. baylyi* ADP1 by natural transformation; Briefly, ADP1 was streaked on LA plate (containing 1% glucose). The cells were incubated at 30^°^C overnight. On the second day, 0.5 μl of plasmid was dropped on the top of a single colony and the cells were incubated for another day. On the third day, the enlarged single colony were picked up and mixed with 50-100 μl LB medium. The mixture was spread onto LA plate (containing 1% glucose and 25 μg/ml chloramphenicol) and incubated at 30 ^°^C until colonies appeared.

The gene *undA* was amplified with primer vs15_2 44 and vs15_1 68 using the genomic DNA of *Pseudomonas putida* KT2440 as template. The fragment was cloned to pBAV1C-chn with BioBrick assembly standard using restriction sites XbaI/SpeI and PstI. The resulting construct was designated as pBAV1C-chn-undA. The gene ‘*tesA* was amplified with tl15 and tl16 from the genome of E. coli MG1655 and cloned to pBAV1C-chn, resulting in pBAV1C-chn-*‘tesA*. The construct pBAV1C-chn-*undA* was further digested with SpeI and PstI and ligated with the previously digested *undA*, resulting in the construct pBAV1C-chn-*‘tesA-undA*. The former two plasmids contain only *undA* or ‘*tesA* respectively. The latter plasmid contains both ‘*tesA* and *undA.*

The transformation of *E. coli* was carried out using electroporation and the transformants was screened on LA plates containing 25 μg/ml chloramphenicol. *A. baylyi* ADP1 was transformed with the three constructed plasmids and the empty pBAV1C-chn. The adapted ADP1 was transformed with pBAV1C-chn-*‘tesA-undA*. The transformation was carried out as described by Metzgar et al. (2004), with an exception that mineral salts medium containing 100 mM ferulate was used as the medium for the transformation. The constructs were verified by restriction analysis.

### 5.6 Cultivation

For testing the functionality of (*ChnR*/*P*_*chnB*_), ADP1 carrying pBAV1C-chn-GFP plasmid was cultivated in minimal salts medium containing 1% glucose, 0.2% casein amino acid and 25 μg/ml chloramphenicol at 30 ^°^C and 300 rpm. When the optical density at wavelength 600 nm (OD600) reached 0.5-1, 1 mM cyclohexanone was added in the culture. Cultivation without the addition of cyclohexanone was used as reference. The cultivation was performed in duplicate. Samples were taken for OD600 and fluorescence after 3 hours of cultivation. For fluorescence measurement, appropriate dilution was made to ensure that samples contained the same amount of biomass. Fluorescence measurement was performed with Spark multimode microplate reader (Tecan, Switzerland) with wavelengths 485 nm (excitation) and 510 nm (emission) and the signal was proportioned to that of the non-induced cells.

The cultivation for plasmid selection was carried out with the three constructed strains, ADP1 *undA*, ADP1 ‘*tesA*, and ADP1 ‘*tesA-undA*. ADP1 containing empty plasmid was used as control. Cultivation was performed in duplicate. Cells were precultivated in 5ml LB medium containing 0.4% glucose and 25 μg/ml chloramphenicol. After overnight cultivation, cells were transferred to 6 ml MA/9 medium containing 5% glucose, 0.2% casein amino acid and 25 μg/ml chloramphenicol and cultivated at 30 ^°^C and 300 rpm. Initial OD was made to 0.05. Cells were induced with 1 mM cyclohexanone after 5 h of cultivation (OD around 1). After 1 h of induction, 5 ml culture was transferred to sealed headspace 20 ml vials (Agilent Technology, Germany) containing a stir bar and incubated at 25 ^°^C and 300 rpm overnight.

The cultivation for 1-undecene production from ferulate was carried out with ADP1 ‘*tesA-undA* and adapted ADP1 ‘*tesA-undA*. Cells were precultivated in mineral salts medium containing 5 mM ferulate and 25 μg/ml chloramphenicol at 30 ^°^C and 300 rpm. The chloramphenicol used in the cultivation was prepared with water to ensure that ferulate is the only carbon source and energy source. ADP1 ‘*tesA-undA* was precultivated for 48 h while adapted strain was precultivated for 24 h. After precultivation, cells were transferred to 110 ml flasks supplemented with 12 ml mineral salts medium containing 110 mM ferulate and 25 μg/ml chloramphenicol. The initial OD was 0.05. Cells were induced after 17 h of cultivation with 1 mM cyclohexanone. After 4.5 h of induction, 10 ml culture was taken from each flask and evenly distributed to two sealed headspace 20 ml vials containing stir bars (each vial contains 5 ml culture). The cells were then incubated at 25 ^°^C and 300 rpm for 20 h.

### 5.7 Analysis methods

The consumption of carbon sources was analyzed with high performance liquid chromatography (HPLC). The samples were collected from the cultures and centrifuged at 20000 g for 5 minutes. The supernatant was taken and filtered with syringe filters (CHROMAFIL^®^ PET, PET-45/25, Macherey-Nagel, Germany). The filtered supernatant was diluted with sterile deionized pure water. The measurement of ferulate concentration was performed with Agilent Technology 1100 Series HPLC (UV/VIS system) equipped with G1313A auto sampler, G1322A degasser, G1311A pump and G1315A DAD. Rezex RFQ-Fast Acid H^+^ (8%) (Phenomenex) was used as the column and placed at 80 ^°^C. Sulfuric acid (0.005 N) was used as the eluent with a pumping rate of 1 ml/minute.

The detection of 1-undecene was performed with SPME-GCMS described by Rui et al. (2014). Briefly, after cultivation, the sealed headspace vials were placed in aluminum block at 25 ^°^C. At the same time, the culture was stirred with a magnetic stirrer. An SPME fiber (d_f_ 30 μm, needle size 24 ga, polydimethylsioxane, Supelco, Sigma-Aldrich) was injected into the vials and held for 12.5 minutes for absorption. GC-MS analysis was performed with Agilent 6890N GC system with 5975B inert XL MSD. The analytes were desorbed from the fiber in a splitless injector at 250 ^°^C for 75 seconds and developed with helium as carrier gas with a flow rate of 1 ml/min. Temperature gradient was applied: 50 ^°^C for 3min, temperature ramped to 130 ^°^C with a rate of 10 ^°^C/min, then ramped to 300^°^C with a rate of 30 ^°^C/min, 300 ^°^C for 5 min.

## 6 Acknowledgements

The research work was supported by Academy of Finland (grants no. 286450, 310135, 310188, and 311986)

